# Oriented Soft DNA Curtains for Single Molecule Imaging

**DOI:** 10.1101/2020.06.15.151662

**Authors:** Aurimas Kopūstas, Šarūnė Ivanovaitė, Tomas Rakickas, Ernesta Pocevičiūtė, Justė Paksaitė, Tautvydas Karvelis, Mindaugas Zaremba, Elena Manakova, Marijonas Tutkus

## Abstract

Over the past twenty years, single-molecule methods have become extremely important for biophysical studies. These methods, in combination with new nanotechnological platforms, can significantly facilitate experimental design and enable faster data acquisition. A nanotechnological platform, which utilizes flow-stretch of immobilized DNA molecules, called DNA Curtains, is one of the best examples of such combinations. Here, we employed new strategies to fabricate a flow-stretch assay of stably immobilized and oriented DNA molecules using protein template-directed assembly. In our assay a protein template patterned on a glass coverslip served for directional assembly of biotinylated DNA molecules. In these arrays, DNA molecules were oriented to one another and maintained extended either by single- or both-ends immobilization to the protein templates. For oriented both-end DNA immobilization we employed heterologous DNA labeling and protein template coverage with the anti-digoxigenin antibody. In contrast to the single-end, both-ends immobilization does not require constant buffer flow for keeping DNAs in an extended configuration, allowing us to study protein-DNA interactions at more controllable reaction conditions. Additionally, we increased immobilization stability of the biotinylated DNA molecules using protein templates fabricated from traptavidin. Finally, we demonstrated that double-tethered Soft DNA Curtains can be used in nucleic acid-interacting protein (e.g. CRISPR-Cas9) binding assay that monitors binding location and position of individual fluorescently labeled proteins on DNA.

## Introduction

Dynamic protein-nucleic acids (NA) interactions play a crucial role in the regulation of many cellular processes. Currently these problems are widely investigated using advanced microscopy-based methods that enable direct monitoring of NA-protein interactions at the single-molecule (SM) level in real time. Information obtained from these experiments is crucially important for building mechanistic models of diverse reactions.^1,2^ Nano- or micro-scopic platforms combined with microscopy techniques become very popular and allow accessing information that is otherwise hidden.^3–5^ However, most of the SM techniques cannot be parallelized and are often technically challenging. Therefore, new high-throughput platforms for SM imaging of protein-NA interactions are in high demand.^6^

One of the best combinations of SM methods with a nanotechnological platform, was the development of the Deoxyribonucleic acid (DNA) Curtains platform. It enabled high-throughput SM imaging by employing nano-engineering, microfluidics, supporting lipid bilayers (SLB) and SM microscopy.^6–8^ This platform utilizes an inert lipid bilayer, which passivates the otherwise sticky surface of the flowcell channel, and mechanical barriers to partition the lipids. Biotinylated DNA molecules that are anchored on the biotinylated lipids via neutravidin (nAv) can be manipulated using hydrodynamic force. Another similar recently developed platform is called DNA skybridge.^9^ It utilizes a structured polydimethylsiloxane (PDMS) surface for DNA immobilization and a thin Gaussian light sheet beam parallel to the immobilized DNA for visualization of DNA and protein interaction at the SM level. The original DNA Curtains platform demonstrated great benefits for studies of many different NA-interacting proteins. However, the original DNA Curtains are less stable and more expensive fabrication-wise than the platform described in this and our previous work.^10^ The skybridge platform contains stably immobilized DNA molecules, but it utilizes rather unusual phenomena for visualization of fluorescently labeled DNA and proteins. However, it is an interesting alternative to the existing DNA Curtains platform.

Recently we demonstrated that streptavidin (sAv) patterns on the modified coverslip surface can be utilized to fabricate biotinylated DNA arrays.^10^ The design of the protein patterns on the modified surface ensures predefined distribution and aligning of the biotinylated DNA molecules on the narrow line-features (> 200 nm). We refer to these aligned molecules as Soft DNA Curtains. The application of hydrodynamic buffer flow allows extension of the immobilized DNA molecules along the surface of the flowcell channel. These Soft DNA Curtains permit simultaneous visualization of hundreds of individual DNA molecules that are aligned with respect to one another and offer parallel data acquisition of diverse biological systems. We showed that Soft DNA Curtains are easy to fabricate in any laboratory having an access to an atomic force microscope (AFM) and objective or prism-based total internal reflection fluorescence microscopy (TIRF).

One of the drawbacks of our previous work was that the double-tethered Soft DNA Curtains had no defined orientation of both-end biotinylated DNA molecules. Such DNA molecules could bind to the sAv line-feature in any direction. Random orientation of DNAs would not create a huge problem because one end of the DNA molecule could be fluorescently labeled and this labeling would allow us to post-orient DNA molecules during data analysis. However, this procedure introduces an extra complication of the experiment.

Here we fabricated the uniformly oriented double-tethered DNA Curtains using heterologous labeling of the DNA molecules by biotin and digoxigenin (dig). We confirmed the defined orientation of DNA molecules using a fluorescent tag introduced asymmetrically to the DNA molecule. These improvements allowed us to demonstrate that double-tethered Soft DNA Curtains can be used in NA-interacting protein binding assay that monitors binding location and position of fluorescently labeled CRISPR-Cas9 proteins on DNA. The well-controlled fabrication procedure of high-quality protein templates was achieved using a portable printing device (PPD) developed especially for this purpose. We increased stability of the immobilized DNA molecules using a more stable alternative to sAv called traptavidin (tAv)^11,12^ as an ink for the fabrication of protein templates.

## Materials and methods

### Chemicals and Materials

Silicone elastomer Sylgard 184 (Dow Corning, Midland, MI, USA) was used for lift-off microcontact printing (μCP) stamp production. For Si master structure production, the gold coated silicon wafers (a 20 nm-thick Au film and a 2 nm Ti adhesion layer, Ssens BV, The Netherlands) were used. Before use, substrates were cleaned in SC-1 solution: ultrapure water, 30% hydrogen peroxide (Carl Roth GmbH, Germany), 25% ammonia solution (Carl Roth GmbH, Germany) at 5:1:1 v/v/v, respectively. Wet chemical etching solution for Au: 20 mM Fe(NO_3_)_3_·9 H_2_O (Fluka, Switzerland), 30 mM thiourea (Fluka, Switzerland) and 1 mM HCl (Sigma-Aldrich, USA) dissolved in ultrapure water saturated with octanol (Sigma-Aldrich, USA). DNA primers were synthesized and purified by Iba-lifesciences (Germany) or Metabion (Germany). Nitrogen gas (purity of 99.999%, ElmeMesser Lit, Lithuania), ultrapure water (Synergy 185 UV, Millipore or Labostar, Siemens), ethanol (99,9%, Merck KGaA, Germany), streptavidin (SERVA, Germany), HEPES (Carl Roth GmbH, Germany), Tris-acetate (Sigma-Aldrich, USA), NaCl (Carl Roth GmbH, Germany), biotin-PEG4-NHS (Jena Bioscience, Germany), Anti-Dig antibodies (Roche). Buffer solutions: A) 33 mM Tris-acetate (pH=7.9, at 25 °C), 66 mM K-Acetate, B) 40 mM Tris, (pH=7.8 at 25 °C), C) 20 mM HEPES (pH=7.5, at 25 °C), 150 mM NaCl.

### Production and purification of proteins

His-tagged tAv was produced and purified according to the published protocol.^11,12^ *E. coli* BL21(DE3) cells were transformed with pET21a tAv plasmid, plated onto Luria-Broth (LB)-Carbenicillin agar plates and incubated at 37 °C overnight. An overnight culture in LB-Ampicillin was grown out of a single colony with shaking 220 r.p.m. and 37 °C. The overnight culture was diluted 100-fold into LB-Ampicillin medium, grown at 37 °C until OD_600_ 0.9, and protein expression was induced with 0.5 mM isopropyl-β-D-thiogalactopyranoside for 4 h at 37 °C. Cells were collected by centrifugation at 5000g and 4 °C for 10 min. The cell pellet was resuspended in a lysis buffer (300 mM NaCl, 50 mM Tris, 5 mM EDTA, 0.8 mg/mL lysozyme, 1% Triton X-100 (pH=7.8, at 25 °C)) and put on a rocker at 80 r.p.m. at room temperature for 20 min. Pulsed sonication of the cell pellet on ice at 30% amplitude was performed afterwards for 10 min. Centrifugation at 27000 g and 4 °C for 15 min followed by washing of the inclusion body pellet in a wash buffer (100 mM NaCl, 50 mM Tris, 0.5% Triton X-100 (pH=7.8, at 25 °C)) was repeated three times. Isolated inclusion bodies were dissolved in 6 M guanidinium hydrochloride (pH=1.5, at 25 °C) and then spun at 17700g and 4 °C for 20 min. Protein precipitation using solid ammonium sulfate was then carried out in order to precipitate tAv from their refolds. The precipitate was resuspended in a minimal volume of PBS at room temperature, centrifuged at 14000g and 4 °C for 5 min and the excess of ammonium sulfate was removed by running the supernatant through a NAP-25 column (GE Healthcare). The tAv was purified using HiTrap chelating column (GE Healthcare) charged with Ni^2+^ equilibrated with a equilibration buffer (300 mM NaCl, 50 mM Tris-hydrochloride (pH=7.8, at 25 °C)). Protein was eluted with elution buffer (300 mM NaCl, 50 mM Tris, 0.5 M Imidazole (pH=7.8, at 25 °C)). The fractions containing tAv were dialyzed into PBS at 4 °C and concentrated by ultrafiltration using a 9 kDa MWCO centrifugal concentrator. The final yield of purification was 3 mg of tAv per 1 liter of initial culture. Wild-type *Streptococcus pyogenes* (Sp) Cas9 was expressed and purified as published previously.^13^

### Production of DNA

Biotinylated oligonucleotides were annealed to the overhang (cos sequences) at either the left, or both ends of bacteriophage λ DNA (48.5 kb, ThermoFisher Scientific). The sequences of the oligonucleotides: 5’-AGGTCGCCGCCC[TEG-digoxigenin]-3’ (right end) and 5’-GGGCGGCGACCT-TEG[Biotin]-3’ (left end) (Metabion). These two oligonucleotides were phosphorylated at the 5’-end using Polynucleotide Kinase (PNK, ThermoFisher Scientific) reaction (1 μM of the respective oligonucleotide, 10 x diluted PNK, PNK buffer, 0.1 mM Adenosine triphosphate, ATP) at 37 °C for 30 min. PNK was inactivated by incubation for 5 min at 95 °C. The λ DNA and the oligonucleotide were mixed at the molar ratio of 1:10, heated to 80 °C, and slowly cooled to room temperature. Subsequently, T4 DNA ligase (ThermoFisher Scientific) was added, and the reaction mixture was incubated at room temperature for 2 h. After the reaction was complete, the DNA ligase was inactivated by heating to 70 °C for 10 min, the excess oligonucleotide was removed using a CHROMA SPIN TE-1000 column (Clontech, USA), and the purified DNA was stored at −20 °C.

For the insertion of an ATTO647N-labeled oligonucleotide complementary to the position 14711 bp from the biotinylated end of the λ DNA we employed the previously described strategy^14^ and followed the more recently described procedure.^15^ 2 μg of λ DNA was incubated for 2 hours with the nicking enzyme Nt.BstNBI (20 units, NEB) at 50 °C in the nickase buffer. The nicked DNA was mixed with a 100-fold excess of 3 oligonucleotides: (5’-pTTCAGAGTCTGA CTTTT[ATTO647N]-3’), (5’-AGGTCGCCGCCC[TEG-digoxigenin]-3’) and (5’-GGGCGGCGACCT-TEG[Biotin]-3’). The mixture was incubated at 55 °C for 20 min and then cooled down at a rate of 0.5 °C/min to 16 °C. Prior to ligation, ATP was added to a final concentration of 1 mM along with 50 units of T4 ligase (ThermoFisher Scientific). The ligation reaction mixture was incubated at room temperature overnight. Any remaining nicking or ligase activity was quenched by adding 20 mM EDTA. The excess oligonucleotides were removed using a CHROMA SPIN TE-1000 column (Clontech, USA), and the purified DNA was stored at −20 °C.

Biotinylated 5 kb long DNA was synthesized by PCR using ΦX174 RF1 DNA (ThermoFisher Scientific) as a template and oligonucleotides 5’ biotin-CGAAGTGGACTGCTGGCGG-3’ and 5’-CGTAAACAAGCAGTAGTAATTCCTGCTTTATCAAG-3’as primers. The product was purified using the GeneJET PCR Purification Kit (ThermoFisher Scientific).

### Fabrication and characterization of a silicon master

Si masters were fabricated and characterized according to the previously published procedure.^10^ Fabrication of the Si master involves formation of a self-assembled monolayer (SAM) from 1-eicosanethiol (HS-C20, AlfaAesar),^16^ surface patterning by the nanoshaving lithography technique using an AFM (NanoWizard3, JPK Instruments AG, Germany) and wet chemical etching. Characterization of the Si master was performed using an upright optical microscope BX51 (Olympus, Japan) and the AFM, operating in AC mode.

### Characterization of printed protein features

The width and morphology of the printed protein features on the glass surface were analyzed with an AFM in buffer C. For that, the glass sample was mounted into the ECCel (JPK Instruments AG, Berlin, Germany) and imaged using the QI-Advanced mode. Before each measurement the probe sensitivity and spring constant were calibrated using the contact-free calibration routine (based on the thermal spectrum of cantilever oscillations) built into the AFM software. The setpoint for measurements was set to 1.5-2 nN tip pushing force.

### Protein nanopatterning by lift-off μCP and the portable printing device

Flat PDMS elastomer stamps for protein lift-off μCP were fabricated according to the previously published procedure.^10^ Briefly, the prepolymer and curing agent (10:1 ratio w/w, Sylgard 184 kit) were thoroughly mixed, degassed in a vacuum desiccator (30 min), poured into a plastic Petri dish and cured in an oven (65 °C for 14 h). The thickness of the cast PDMS elastomer was ~2 mm. The PDMS surface that was in contact with the Petri dish was treated as the flat one.

The lift-off μCP was performed similarly to the published procedure.^10,20^ The PDMS elastomer (5 x 5 mm^2^ dimensions) was immersed in isopropanol for 10 min, held by tweezers and dried for 15 s, placed on a plastic Petri dish and dried for another 10 min. The Si master was immersed in isopropanol (20 min), held by tweezers and dried, and cleaned by air plasma (5 min, ~500 mTorr, high-power mode, PDC-002, Harrick, USA). To homogeneously cover the PDMS surface with a film of the protein ink the PDMS stamp was placed in a clean plastic Petri dish with its flat side facing up. Then a 60 μL drop of specified protein solution (in buffer A) was placed on the PDMS surface, mixed with the tip of pipette, and kept for 10 min. After incubation the protein ink was removed from the PDMS stamp by sucking it out with the pipette tip. Then, it was held with tweezers and washed with 5 mL of buffer A using a 1 mL pipette, ~50 mL ultrapure water using a wash bottle, and dried under N_2_ gas stream.

For the printing procedure we build a semi-automated printing machine, which allowed us to apply different printing pressure (PP, see SI file for the detailed description). First, the cleaned Si master was placed on the silicon rubber on the bottom of the printing machine (SI Fig. 2A). The dried PDMS stamp was placed facing flat side-up on a piece of glass (10 x 10 mm^2^ dimensions), which was covered with the double-sided sticky tape (SI Fig. 2B). Next, the PDMS stamp on the piece of glass was placed on the Si master using tweezers. The syringe holder was mounted on the top of the device and the pressure (PP=0.6 mL, unless stated otherwise) was applied to the PDMS stamp on the Si master using the distant syringe (SI Fig. 2C). The contact in between the patterned Si surface and PDMS stamp was established for ~15 s, then the distant syringe was released and the glass slide with PDMS elastomer was removed from the Si master using tweezers. Subsequently, the silanized and PEGylated (methoxy-PEG-SVA and biotin-PEG-SVA, both 5 kDa, Lyasan Bio, USA) glass coverslip (25 x 25 mm^2^, #1.5, Menzel Glaser) was placed instead of the Si master using tweezers. The PDMS stamp was transferred onto the PEGylated glass coverslip and kept for 1 minute under the pressure (PP=0.6 mL, unless stated otherwise) applied by the distant syringe. The surface of the glass coverslips was modified in the same way as described previously,^21^ using the biotin-PEG:methoxy-PEG (bt-PEG:m-PEG) ratio 1:10 (w/w). In order to minimize non-specific protein adsorption to the surface, we performed a second round of PEGylation with the short NHS-ester PEG molecules (333 Da) according to the published procedure.^22^ Next, the glass slide with the PDMS stamp was removed from the glass coverslip using the tweezers and discarded. The patterned glass coverslip was assembled into the flowcell, which was prepared as described earlier.^10^ The Si masters were reused for the lift-off μCP multiple times and in between the experiments they were stored in 100% isopropanol solution.

### TIRF microscopy

The employed home-build TIRF microscopy setup was described previously.^10^ This microscopy setup was equipped with three different wavelength lasers: 488 nm, 532 nm and 635 nm (all 20 mW, Crystalaser, USA). These combined beams were directed to the objective (100x, 1.4NA, Nikon) using a quad-line dichroic mirror (zt405/488/532/640rpc, Chroma Technology Corp), which was placed in the upper filter cube turret installed in the microscope body (Nikon Eclipse Ti-U). The laser power before the objective was set to 2.5 mW for both 532 nm and 632 nm lasers, and to 0.1 mW for the 488 nm laser, respectively. The exposure time of the EMCCD camera (Ixon3, Andor) was set to 100 ms. The microscopy images represented in the article and in the SI file were averaged over 10 consecutive frames, thus improving the signal-to-noise ratio (SNR). The penetration depth of the evanescent field was set to ~300 nm for all wavelengths of excitation. This setup was equipped with a custom-build feed-back control system to compensate the Z-axis drift of the sample and keep it stably in focus. The average line quality factor was calculated using the formula described in our previous publication.^10^

### DNA immobilization

First, the channel of the flowcell was filled with buffer B. To enhance surface passivation against non-specific protein adsorption, we injected 5% Tween-20 solution in buffer B into the channel of the flowcell, incubated for 10 min and washed out with 600 μl of buffer A.^23^ Next, the biotinylated DNA (~30 pM, in buffer B) was added and incubated for at least 15 min. The excess of unbound DNA was washed out with ~300 μl of buffer A. Then DNA was labeled with the DNA intercalating green fluorescent dye – SYTOX green (SG, ThermoFisher Scientific, USA) at a concentration of ~0.4 pM (in imaging buffer: buffer A supplemented with 0.2% Tween-20, 1 mM Dithiothreitol, DTT). The SG dye was present during the entire time of the experiment. In the case of the second DNA end tethering, the close-loop circulation was employed and 5 μl of biotin-anti-dig (bt-anti-dig) antibody was added (this resulted in ~0.05 mg/mL concentration) and incubated for at least 10 min at low speed (~0.1 mL/min). After 10 min, the speed of the buffer flow was increased to ~1 mL/min and kept constant for 20 min. Then, to remove the excess of unbound bt-anti-dig the flowcell was washed with 500 μL of buffer A in the open-loop circulation. Finally, 100 μl of imaging buffer was injected into the flowcell in order to reveal bound DNA.

### Cas9 labeling and complex assembly

SpCas9 complex was labeled using ATTO647N conjugated oligonucleotide, which was hybridized with the tracrRNA 5’-end in the crRNA:tracrRNA duplex. Briefly, the tracrRNA 3’-modified for hybridization with the ATTO647N labeled oligonucleotide was obtained using PCR with 5’-TAATACGACTCACTATAGGGCAAAACAGCATAGCAAGTTAAAATAAGG-3’ and 5’-GCGCACGAGCAAAAAGCACCGACTCGGTGCC-3’ primers from the pUC18 plasmid containing RNA encoding sequence followed by *in vitro* transcription (TranscriptAid T7 High Yield Transcription Kit, Thermo Fisher Scientific) and purification (GeneJET RNA Purification Kit, Thermo Fisher Scientific). Next, the oligonucleotide 5’-TTGCGCACGAGCAAA-3’ (Metabion International AG) which is complementary to the tracrRNA 5’-end was labeled with ATTO647N-NHS (1:60 DNA:dye molar ratio) and purified using G-25 micro spin column (Illustra, GE Healthcare). The measured labeling efficiency of ATTO647N-oligonucleotide was >80%. Subsequently, the assembly of crRNA:tracrRNA duplex and the hybridization of tracrRNA with ATTO647N-oligonucleotide were performed simultaneously by mixing the equimolar amounts of synthetic crRNA (Synthego), which contains 5’-GAAATCCACTGAAAGCACAG-3’ target site, tracrRNA and ATTO647N-oligonucleotide along with 5x annealing buffer (Synthego), then heating the mixture to 80 °C and allowing it to slowly cool down to room temperature. Finally, Cas9-RNA complex was assembled from SpCas9 and crRNA:tracrRNA-ATTO647N (1:2 protein:RNA molar ratio) in reaction buffer (10 mM Tris-HCl, pH=7.5, at 37 °C, 100 mM NaCl, 1 mM EDTA, 0.5 mg/ mL BSA, 1 mM DTT) at 50 nM final concentration at 37 °C for 30 min. The complex was diluted to the concentration 0.2 nM in the imaging buffer and injected into the flowcell for TIRF microscopy.

### Cas9 binding location and duration characterization

Binding location characterization was done using a custom-written automated procedure. It fits each Cas9-ATTO647N complex (red-fluorescent spot in TIRF images acquired under 635 nm wavelength excitation) to the 2D Gaussian function with the help of the detection of clusters of interconnected pixels that have values above the manually-set threshold. This protocol is similar to the previously published one.^24^ All procedures were written using Igor Pro (Wavemetrics, Inc.) and are available upon direct request to the authors. Both center coordinates (x and y) were recorded for each detected fluorescent spot that had fitting error of all parameters was <60% from the parameter value. We mainly detected individual stable binding events of various durations without any diffusion characteristics. After fitting, we manually examined the fitted data and selected algorithm-suggested interconnected fit points (stable binding states) that contained more than 5 points (duration at least 0.5 s) in them. This analysis allowed us to extract Cas9 binding state durations (i.e. dwell times) and correlate them with the position on the DNA substrate.

## Results and discussion

To utilize the DNA Curtains platform for complex protein-NA interaction studies, it is required to obtain double-tethered DNA molecule arrays with defined orientation. Therefore, in this work, we upgraded the existing Soft DNA curtains platform^10^ and further optimized its fabrication method by introducing several new steps that made the platform more stable and more controllable.

### Optimization of DNA arrays fabrication

Here, we expanded our previous work^10^ and demonstrated that lift-off μCP patterned sAv or, as we show here later, tAv protein templates (Fig. 1A) can be employed for the self-assembly of biotin labeled DNA molecules in the flowcell on the glass surface (Fig. 1B). In principle, the design of the template ensures the distribution of biotinylated DNA molecules on predefined narrow protein line-features fabricated on the modified glass coverslip, which is otherwise resistant to nonspecific protein interactions. Application of the buffer flow pushes the DNA molecules through the flowcell channel while their biotinylated ends remain tethered. Also, the design of the template is such that the line-spacing distances of templates are sufficiently long and avoid overlapping of DNA molecules immobilized on the neighboring line-features. Such protein array templates can be considered as a soft functional element and therefore we term our platform the Soft DNA Curtains.

**Figure 1:**
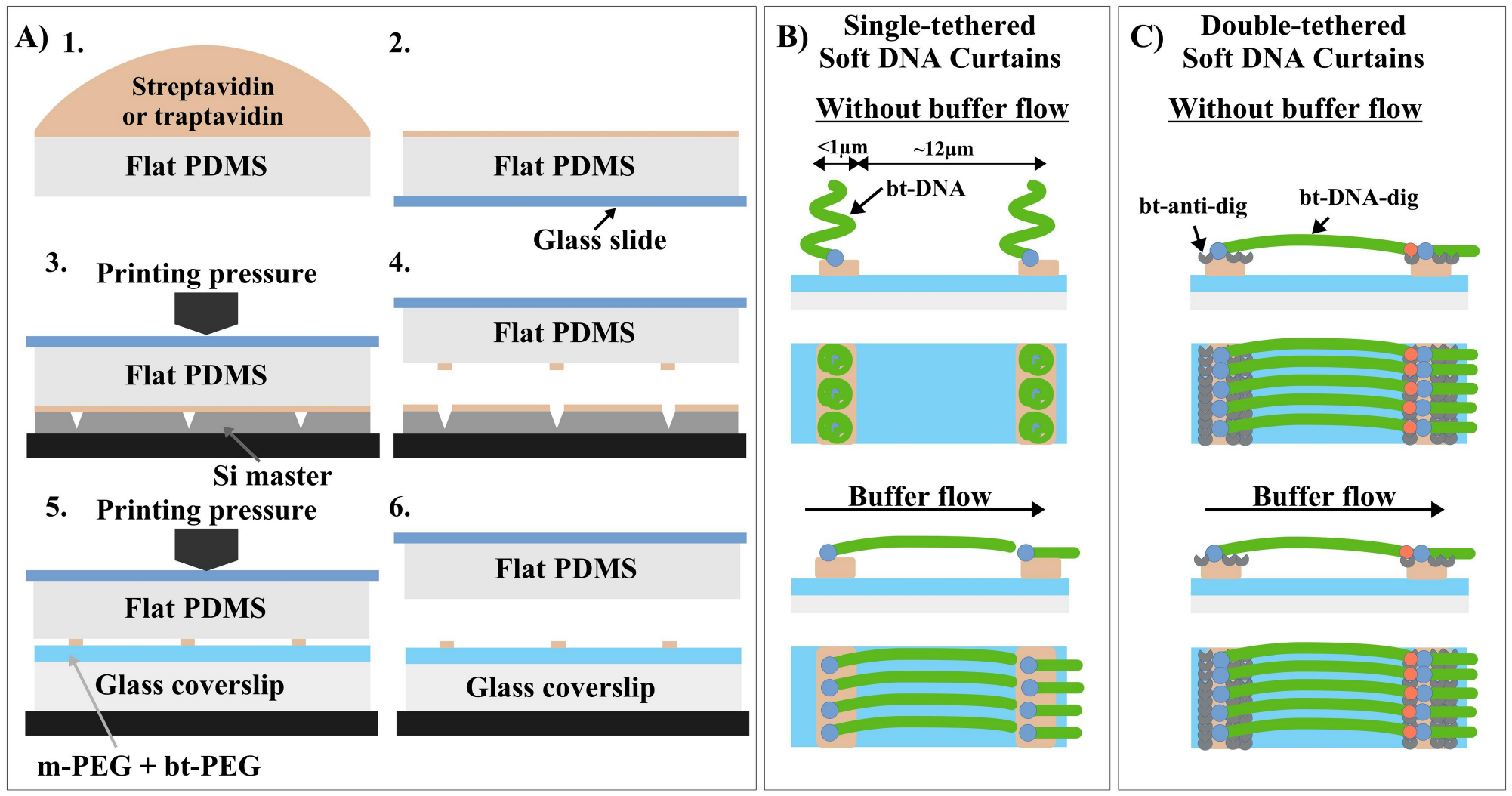
Diagram explaining the main steps of Soft DNA Curtains fabrication. **A)** Diagram of protein liftoff micro-contact printing (μCP): (1) inking of a planar PDMS stamp with streptavidin (sAv) or traptavidin (tAv) ink; (2) drying of PDMS under a stream of N2 gas; (3–4) selective subtraction of sAv (or tAv) by contacting the Si master with the inked PDMS under pressure applied by the Portable Printing Device (PPD) followed by a lift-off; (5–6) μCP under pressure applied by the PPD of sAv or tAv onto a glass coverslip modified with methoxy- and biotin-PEG mixture. **B)** Diagram of single-tethered Soft DNA Curtains illustrating immobilization and alignment of biotin (bt) labeled DNA on the fabricated sAv (or tAv) templates. **C)** Diagram of double-tethered Soft DNA Curtains illustrating immobilization and alignment of bt and digoxigenin (dig) labeled DNA on the fabricated sAv (or tAv) templates via biotin-sAv/tAv and dig-anti-dig interactions.

First, to achieve protein array templates allowing desired distribution of biotinylated DNA molecules, we fabricated Si masters using the previously described procedure^10^ with line-widths ranging from ~200 nm to 1 μm and line-spacing corresponding to ~75% (~12 μm) of the mean extension of λ DNA.^6^ The dimensions of Si masters’ patterned area were from 0.5 × 0.9 mm^2^ to 2.5 × 1.2 mm^2^. Typical line-depth of the Si masters used in this work is ~200 nm. SI Figure 1 shows the Si masters’ overall optical images, line-width and -depth measurements using AFM. SI Table 1 summarizes the characteristics of Si masters that were measured by AFM.

To improve the patterning reproducibility and to control the PP in the lift-off μCP, we built the PPD, which is similar to the previously published device.^25^ However, our PPD was assembled from commonly used parts in an optics laboratory and does not require sticking of the PDMS stamp to the moving piston (SI Fig. 2). In addition to that, our PPD employs PDMS stamp attachment to the glass slide surface, which helps to keep the stamp flat (Fig. 1A and SI Fig. 2B). It is worth noting that a similar effect (PP vs. protein array quality) could be achieved by changing the lift-off μCP printing time, but that would tremendously increase its duration. In our experiments the pressure applied by this easy to use and relatively simple device ranged from ~7 N/cm^2^ to ~13.5 N/cm^2^. To keep it simple, instead of N/cm^2^ we chose to report the PP in terms of the position of the syringe 3 piston (SI Fig. 2C). We calibrated this value and the results are given in SI Figure 3. To test the quality of sAv line-features printed using PPD on the m-PEG/bt-PEG (10:1 w/w) modified glass coverslip surface, we immobilized 5 kb long biotinylated double-stranded DNA (dsDNA) (Fig. 2A). TIRF images showed DNA molecules mainly immobilized on the sAv line-features, but their density was dependent on the applied PP (Fig. 2B and SI Fig. 4). Quantitatively the best results were obtained with a PP of 0.6 mL (SI Table 2). We noticed that at 0.45 mL and especially at 0.3 mL PP DNA immobilization on the line-features was poor and lines became discontinuous. This could be the result of either PDMS touching the bottom of the inscribed lines in the Si master during the printing procedure or pressure-induced inactivation of sAv. To test the first possibility, we performed AFM imaging of sAv lines printed with distinct pressures on the modified coverslip. Results of these measurements showed no evidence of either line breaks or line-width change (Fig. 2B). Therefore, we concluded that too high PP (starting at 0.45 mL) inactivates a fraction of sAv on the surface, which results in reduced binding of biotinylated DNA.

**Figure 2:**
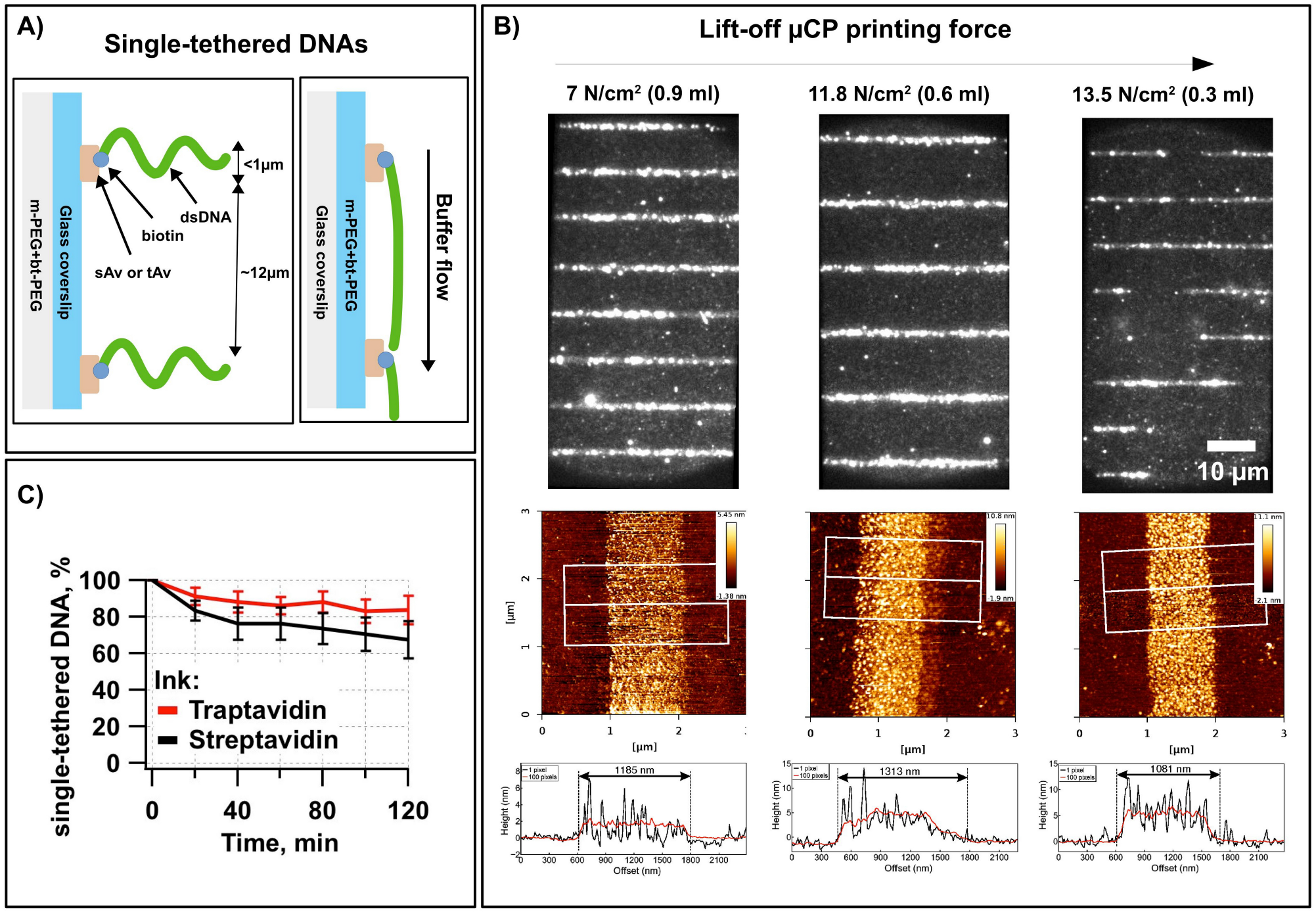
Optimization of DNA arrays fabrication. **A)** Schematic of the single-tethered Soft DNA Curtains design shows PEG monolayer on a glass coverslip and printed streptavidin (sAv) or traptavidin (tAv) line features, which enables specific one-end immobilization of the biotinylated λ DNA (48.5 kb) molecules. DNA molecules are tethered to the line features and responds to a hydrodynamic force by extending parallel to the surface. **B)** Effect of printing pressure (PP) on the quality of short DNA molecule arrays. Top panel shows TIRF images of 5 kb long biotinylated DNA molecules stained with SYTOX green (SG), which were immobilized on the sAv line-features fabricated on modified coverslip. PP is indicated above each image. Bottom panel shows AFM images and their line-profiles (1 and 100 pixels) of the sAv line features fabricated on the modified glass coverslip at the same pressure as the TIRF images. [sAv] = 0.02 mg/mL, Si master #1. **C)** Stability test of single-tethered Soft DNA Curtains – λ DNA molecules immobilized on a sAv (or tAv) array template and stained with SG. Images were acquired every 20 min for a period of 2 h. In between acquisitions, there was no buffer flow applied. During the acquisition, 20 frames were acquired at buffer flow of 1 mL/min and 20 frames without the flow. Graph shows the average number of single-tethered DNA molecules that extended to the full length. Average was taken over line-features and the error bars represents SD. Si master #3.

Another parameter that we assessed in order to optimize the immobilization of biotinylated DNA molecules was the sAv concentration during PDMS elastomer inking under constant PP of 0.6 mL.

In these experiments, the sAv ink concentration ranged from 0.013 to 0.027 mg/mL. We performed lift-off μCP with sAv ink, assembled the flowcell, and immobilized the biotinylated 5 kb long DNA molecules. Acquired TIRF images showed similar results as previously observed^10^ – the highest quality protein templates were fabricated at the moderate sAv concentration of 0.017 mg/mL (SI Fig. 5 and SI Table 2). The optimal range of sAv concentration was rather narrow, since concentrations 52% higher or lower than optimal concentration immediately gave worse results. The obtained optimal sAv ink concentration under 0.6 mL PP is similar to the optimal sAv concentration without applied PP.^10^ However, in contrast to the manual lift-off μCP performed by hand without application of the pressure, the PPD device allows production of a consistent and high-quality protein template across the entire patterned area (SI Fig. 6). This is the main advantage of lift-off μCP using PPD in comparison to the manual procedure.

In our previous work, we showed that the number of single-end tethered biotinylated λ DNA molecules decreased slowly over time, with a half-life of > 2 h,^10^ in a good agreement with the expectations for a high-affinity biotin-sAv interaction. However, a recently developed super-stable variant of sAv – called traptavidin (tAv)^11^ – should allow us to observe immobilized biotinylated DNA molecules for an even longer period of time. We verified our tAv functionality (see SI file and SI Fig. 7) and then tested whether tAv is suitable for fabrication of the fixed DNA molecule arrays. For these experiments we used the lift-off μCP with variable tAv concentration, which ranged from 0.015 mg/mL to 0.06 mg/mL, at constant PP of 0.6 mL. Once line-features were formed and the flowcell was assembled, we immobilized biotinylated 5 kb long DNA molecules. TIRF images showed that the optimal tAv concentration was ~0.03 – 0.02 mg/mL (SI Fig. 8C-D). Both at lower and higher concentration than the optimal we observed more DNA bound in the interline areas or lower DNA density on line-features (SI Fig. 8A-B and E). This visual inspection was also well reflected by the quantitative QF-based characterization (SI Table 2). However, the absolute density of DNA molecules immobilized on the line-features seems to be lower than that with sAv, but this could be rationalized by the different activity of tAv, which requires a higher concentration of DNA molecules to achieve similar densities. Here we decided to use the same concentration of sAv and tAv for the sake of consistency.

To test whether tAv indeed allows observing the immobilized DNA molecules for a longer period of time than sAv, we fabricated single-tethered Soft λ DNA Curtains on sAv (0.017 mg/mL) and tAv (0.03 mg/mL) line-features at a constant PP of 0.6 mL. Next, we performed the stability test of bound DNA molecules by performing the acquisition cycles of image series every 20 minutes. During these cycles 20 frames were acquired with the 1 mL/min buffer flow and 20 frames without the flow. Between the acquisitions there was no buffer flow applied. Acquired TIRF images showed that the number of full-length single-tethered DNA molecules anchored to the surface decreased slowly over time, with the half-life > 2 h for both sAv and tAv (SI Fig. 9). DNA molecules were dissociating slower from tAv than from sAv, and that is in good agreement with the expectation for the lower biotin dissociation constant of tAv.^11^ Namely, after 2 h of observation time only ~20% of DNA molecules were dissociated from tAv line-features, while ~40% of them were dissociated from sAv (Fig. 2C). These results proved tAv to be better suited for the long-lasting experiments, which require observation of the same DNA molecules, and can be more beneficial for all types of DNA Curtains.

### Assembly and characterization of double-tethered Soft DNA Curtains

The patterns of our platform utilize tAv functional elements, and an overview of the general design is presented in Figure 1C. The biotinylated DNA end is first tethered on the protein line-features on the modified glass coverslip surface in the flowcell. In the absence of the buffer flow (hydrodynamic force), the molecules are distributed on the line-features, but lie outside the penetration depth of the evanescent field (Fig. 3A). Application of the buffer flow pushes the DNA through the flowcell channel while biotinylated DNA ends remain tethered. The other end of λ DNA was modified with the dig in order to tether it onto the neighboring protein line-feature. The line-spacing distance was optimized for the length of the λ DNA. The line-features themselves are designed to represent a sufficiently large surface, that can be coated by bt-anti-dig (for functionality verification of bt-anti-dig see SI file and SI Fig. 10). When biotinylated DNA molecules are immobilized on the line-features, we apply slow flow of the buffer containing bt-anti-dig (Fig. 3B). This step allows us to coat the protein line-features with the bt-anti-dig. At the next step the immobilized DNA molecules are stretched by increasing the buffer flow rate. The dig-modified ends of DNA molecules should bind the antibody-coated line-features. This strategy allows us to hold DNA molecules stretched parallel to the surface even when no buffer flow is applied in the flowcell (Fig. 3C).

**Figure 3:**
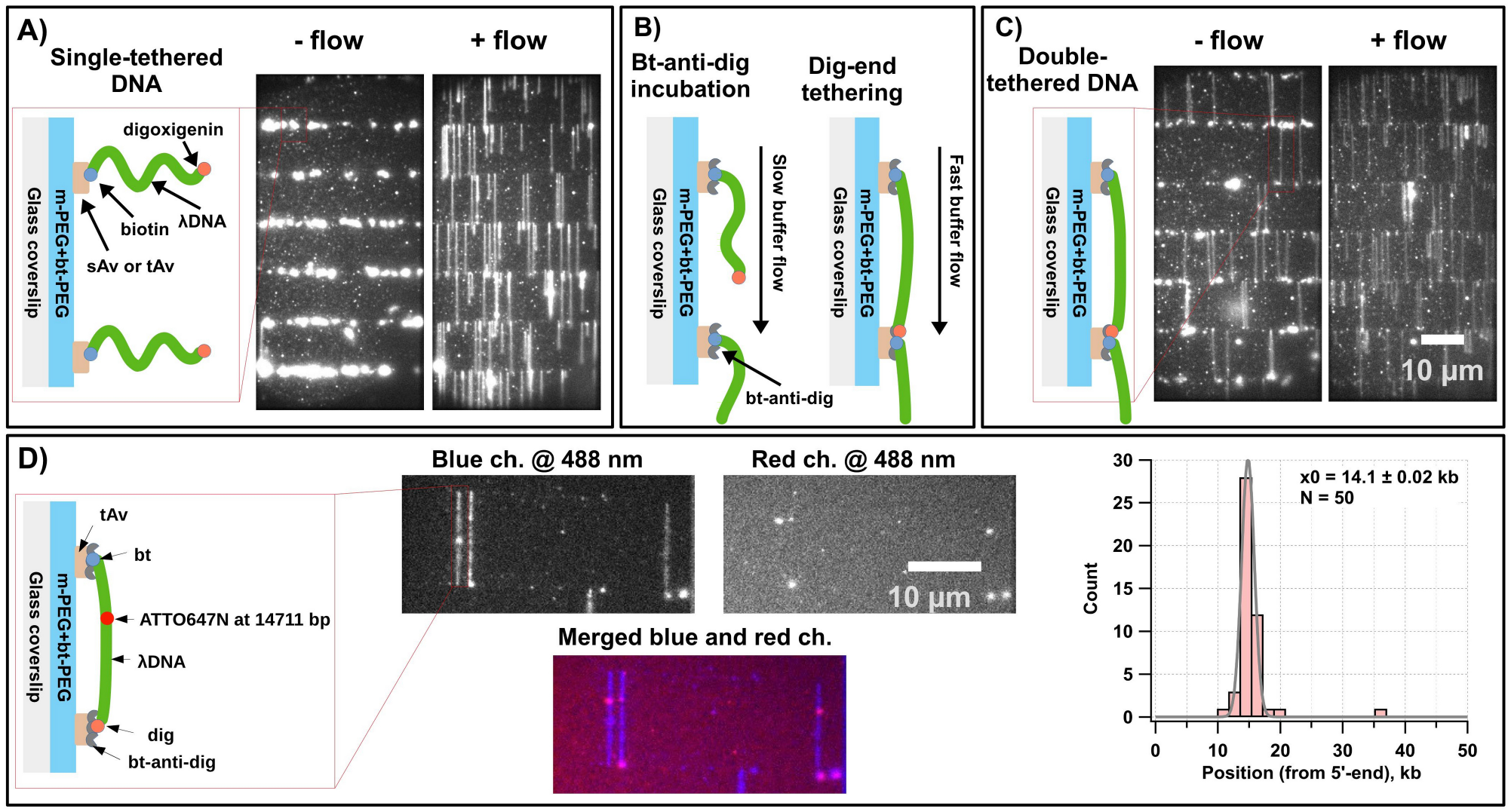
Double-tethered Soft DNA Curtains with defined orientation. **A)** Cartoon illustrates the printed traptavidin (tAv) line-features that enable specific one-end immobilization of the biotin-λ DNA-digoxigenin (bt-λ DNA-dig, 48.5 kb) molecules. TIRF images shows that in the absence of the buffer flow (-flow), DNA molecules are aligned to the line-features. They are responding to a hydrodynamic force (+flow) by extending parallel to the surface. **B)** To tether the dig-labeled end of bt-λ DNA-dig, continuous slow flow of buffer containing biotinylated anti-dig (bt-anti-dig) antibody is applied and DNA molecules are dragged slightly, but do not reach the neighboring line-feature. At the next step, buffer flow rate is increased and the dig-labeled DNA ends encounter the neighboring line-feature, which is now covered with the bt-anti-dig, and the dig-labeled ends become anchored through dig – anti-dig interaction. **C)** Cartoon illustrates the double-tethered bt-λ DNA-dig molecule after dig-labeled end tethering. TIRF images shows that those DNA molecules that remain stretched without buffer flow (-flow) were successfully both-end tethered. **D)** Cartoon illustrates the DNA immobilization strategy and internal ATTO647N tag, which was located at 14711 bp from the biotinylated DNA end. TIRF images shows SG stained DNA molecules in the absence of buffer flow. Excitation wavelength and emission channel is indicated above each image. Histogram showing the distribution of ATTO647N locations that were determined by fitting the images to 2D Gaussian functions. Si master #8.

Line-feature width of the tAv patterns was assessed in order to optimize assembly of the double-tethered Soft DNA Curtains. Three different patterns of tAv were fabricated on the separate coverslips using Si masters #4, #5 or #6 at PP of 0.6 mL. The double-tethered Soft DNA Curtains were assembled on the patterned coverslips as described above (Fig. 3). The anchoring efficiency was tested for tAv patterns made with variable line-widths, but constant line-spacing (i.e. ~12 μm). As expected, the wider lines allowed us to achieve more efficient anchoring. Percentages and densities (average number of both-end anchored DNA molecules per line-feature) of both-end anchored DNA molecules are presented in SI Table 3. Approximately 79% of the anchored DNA were double-tethered with 1 μm wide line-features, 74% with 500 nm and 71% with 350 nm wide line-features. More apparent differences were observed in densities of double-tethered DNA molecules. For the 1 μm-wide line-features it was ~4.9, for 500 nm features ~3.6 and for 350 nm features ~1.6 double-tethered DNA molecules per line-feature. Thus, the thicker the line-feature, the higher the density of both-end tethered DNA molecules.

Once line-feature width was optimized, line-separation distance of the tAv patterns was assessed. Four different patterns of tAv were fabricated on the separate coverslips using Si masters #5, #7, #8 and #9 at PP of 0.6 mL. The double-tethered Soft DNA Curtains were assembled on the patterned coverslips as described above (Fig. 3). This time, the anchoring efficiency was tested for tAv patterns made with variable line-separation distances, but constant line-widths (i.e. ~0.5 μm). As expected, the most efficient anchoring occurred with the 12 and 13 μm line-spacing distances. Percentages of both-end anchored DNA molecules are presented in SI Table 4. Approximately 80% of the anchored DNA were double-tethered with 13 μm, 74% with 12 μm, 48% with 11 μm, and 20% with 14 μm separation distance line-features.

When line-width and -separation distance were optimized, we tested the orientation of λ DNA fragments on the double-tethered Soft DNA Curtains using a fluorescent tag (ATTO647N at the position 14711 bp) introduced asymmetrically to the specific position of the λ DNA. As mentioned above, we used differential chemistries (biotin on the left end, and dig – right end) to tag two ends of the DNA (Fig. 3C). Therefore molecules within the double tethered curtains should be immobilized in a defined orientation. To confirm that the DNA was oriented correctly, we assembled double-tethered Soft DNA Curtains from the ATTO647N labeled λ DNA (bt-λ DNA ATTO647N-dig) as described above (Fig. 3C). The ATTO647N labels were present at a single location within the DNA molecules and aligned with one another. Their mean position was found to be 14.1 ± 0.02 kb (N = 50) from the biotinylated DNA end. This result coincided well with the expected location and practically no ATTO647N labels were observed at other locations.

### Deployment of double-tethered Soft DNA Curtains for visualization of protein-nucleic acid interactions

To demonstrate that double-tethered Soft DNA Curtains can be utilized to visualize protein-DNA interaction, we selected previously characterized Cas9 nuclease from the CRISPR-Cas system of *S. pyogenes* (Sp), that is involved in bacterial defense against foreign invading DNA and has been adopted as a genome editing tool.^26^ Soft DNA Curtains were assembled from bt-λ DNA-dig molecules and tAv line-features covered with the bt-anti-dig. The Si master #8 was used to fabricate the tAv line-features on a PEGylated coverslip at PP of 0.6 mL. Cas9 complex targeting double-tethered λ DNA 31.3 kb from the biotin-labeled end was fluorescently labeled with ATTO647N-oligonucleotide that was complementary to the tracrRNA 5’-end (Fig. 4A). The SpCas9-ATTO647N complexes were diluted to 0.2 nM concentration in a Mg^2+^-free buffer and injected into a flowcell.

**Figure 4:**
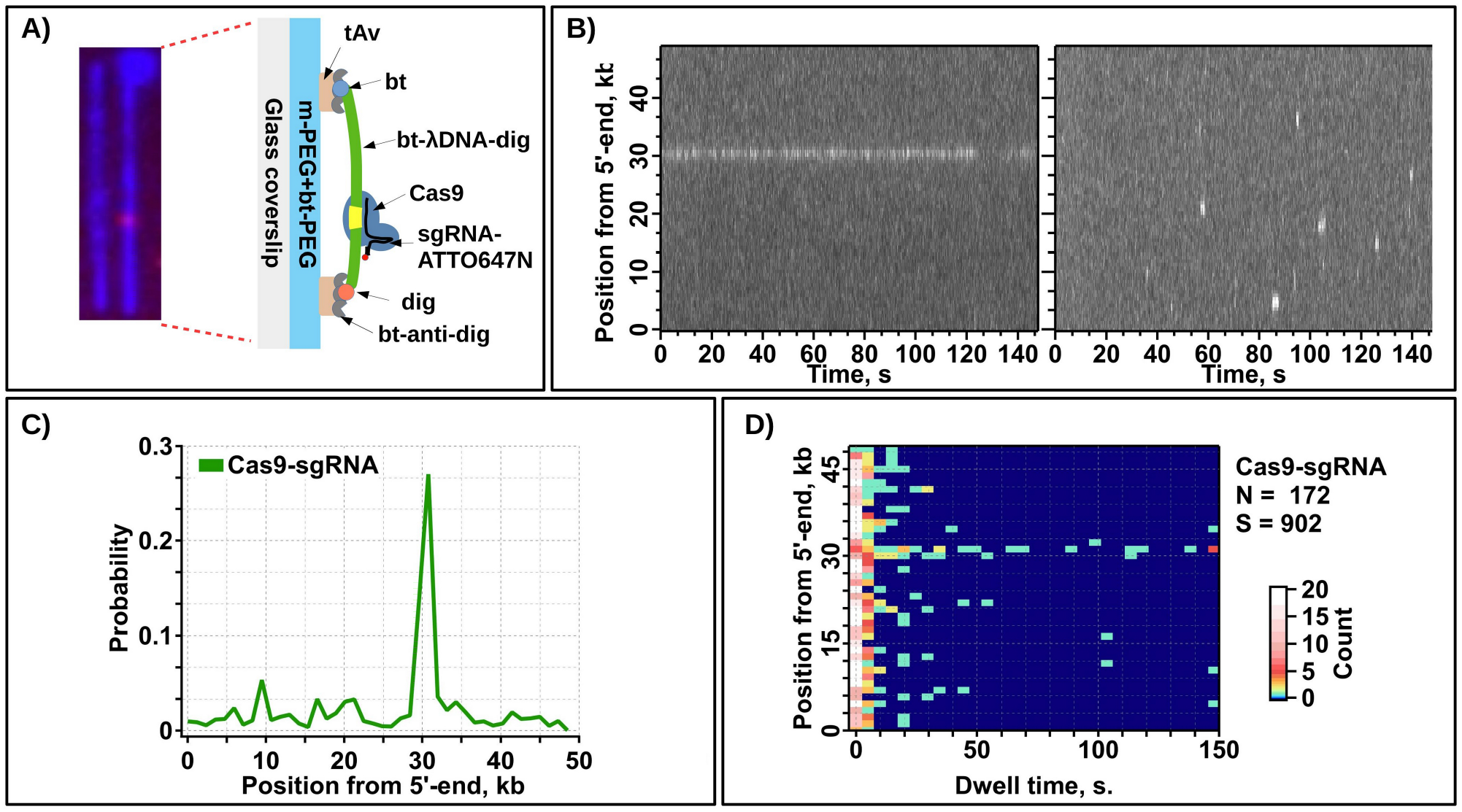
Double-tethered Soft DNA Curtains assay for binding location studies of SpCas9 protein. **A)** Merged blue and red channel TIRF images representing SG-stained λ DNA (blue) with bound SpCas9 (red). Schematic representation of the assay depicts SpCas9, which was programmed with ATTO647N-labeled crRNA-tracrRNA (Cas9-ATTO647N) targeting site of λ DNA located at 31.1 kb from the biotinylated DNA end. **B)** Representative kymograms made from individual DNA molecules. **C)** Histogram of SpCas9-ATTO647N binding events distributions determined by 2D Gaussian fitting. **D)** SpCas9-ATTO647N binding position vs. dwell time 2D histogram plot. The plot was made from 172 DNA molecules and contains 902 individual binding events. Color code represents the counts.

TIRF images show an example of double-tethered curtains with bound Cas9-ATTO647N, where DNA is colored blue, and the protein is colored red (Fig. 4A). This image represents a single 100 ms long image taken from 150 s video (the full length video is not shown). Two representative kymograms extracted from two different DNA fragments show long-lasting binding events occurring on the target site (31.3 kb) and short binding events occurring on the non-target site (Fig. 4B left and right, respectively). A binding profile of Cas9-ATTO647N was obtained from individual DNA molecules (Fig. 4C). There were 172 DNA molecules and 902 binding events of proteins in total monitored during this experiment. As in Figure 4, the DNA-bound Cas9-ATTO647N demonstrated broad binding distribution along the full length of the λ DNA. However, in total protein spend more time at the target site (Fig. 4C). There were more binding events on the left side of the DNA, which likely is dictated by higher GC content. Binding events on target had significantly longer dwell times (some of them lasted as long as the acquisition time 150 s) than on non-target binding, which lasted < 20 s (Fig. 4D). On average target binding events lasted for ~52 s and non-target binding events ~7 s (SI Fig. 11). We note, that this experiment was conducted in the absence of Mg^2+^ ions in the imaging buffer. Therefore, obtained results suggest that Mg^2+^ ions are not essential for specificity of SpCas9 target recognition. This experiment provides direct evidence that the double-tethered Soft DNA curtains can be utilized to visualize the DNA and fluorescently labeled proteins’ interaction at the SM level.

## Conclusion

The oriented double-tethered DNA Curtains allow intuitive and simple parallel examination of hundreds of SM in a single TIRF experiment. In this work, we showed that it is possible to fabricate oriented and aligned Soft DNA Curtains using high-quality protein template-directed assembly of biotinylated DNA molecules and hydrodynamic force. Also, we showed by the Cas9 binding experiment that our oriented double-tethered Soft DNA Curtains are suitable for visualization and characterization of individual NA-interacting protein studies. We believe that each of the optimizations and improvements described in this work will be useful for SM studies and make the Soft DNA Curtains platform better characterized.

## Acknowledgments

This study was funded by Research Council of Lithuania [S-MIP-17-59 for E.M. and M.T. and Dotsut-611 for A.K.]. We are grateful to Dr. R. Valiokas for valuable advice and discussion throughout this work. We thank Arunas Silanskas for purification of Sp Cas9 protein.

