## Supplemental information for "Oriented Soft DNA Curtains for Single Molecule Imaging"

**Portable printing device (PPD).** The basic principles of our PPD device (SI Fig. 2) are similar to the previously published one,[24] and it was assembled from commonly used parts in an optics laboratory. As the SI Fig. 2A shows, firstly we placed the Si master on a silicone rubber sheet attached to a glass slide (microscope slide, 1 mm thickness, Heinz Herenz Hamburg), which was glued to a base plate (3BP4-02, Standa) using a piece of double sided sticky tape (9088-200, 3M). We made holes in the centers of the silicone rubber sheet, the glass slide, and the double-sided sticky tape. To keep the Si master fixed on the silicone rubber sheet, we sucked out the air through this hole. For this purpose, we glued a nanoport (10-32, IDEX Health & Science) to the center of the under side of the glass slide and connected it to a 2 mL syringe (syringe 1 in the SI Fig. 2A) via a 30 cm long tubing (a suitable tubing clamp was installed to clamp the tube after the air was sucked out). After this step, we attached a PDMS elastomer with a protein ink monolayer flat-side-up onto the glass slide using double-sided sticky tape (SI Fig. 2B). Next, we held the glass slide with tweezer and positioned the PDMS elastomer facing inked-side-down onto the Si master. To apply printing pressure, we made a system of two tubing-connected syringes (SI Fig. 2C, 1 mL Omnifix-F, Braun). First, we fitted the first syringe (syringe 2 in the SI Fig. 2C) into the center hole of two connected base plates (3BP4-02, Standa). Second, we filled the syringes and the tubing with tap water and fixed the syringes to the 15 cm long tubing (228-0703, VWR) using tube clamps (EX52.1, Carl-Roth) and 3 cm long adaptor tubings (N874.1, Carl-Roth). These adaptor tubings were placed on the ends of the 15 cm long tubing and made the contact with the syringe stronger. Once the system was assembled, we mounted it onto mounting posts (3MP-25+3AH6-4, Standa) using suitable screws (M6, 3 cm long). Finally, we applied the printing pressure to the PDMS elastomer using the second syringe (syringe 3 in the SI Fig. 2C) and controlled the printing pressure by visually inspecting the volume scale on the second syringe.

**Printing force measurements.** To measure the actual printing force applied with the device, we employed a force sensitive resistor (FSR 400 Series Round, 5.08 $mm^2$ active area, 0.35 mm nominal thickness, Interlink Electronics, Inc.), which was placed in between syringe 2 and the glass slide holding the PDMS elastomer (SI Fig. 2C). One leg of the FSR device was connected to the +5V output port of the Arduino Uno card and the other leg was tied to a measuring resistor (10 kΩ) in a voltage divider configuration (i.e. one end was connected to ground and the other to the analog input port of the Arduino Uno card). The output voltage ($V_{out}$), which was monitored using the Arduino Uno card increased with increasing force. To calculate the actual force in Newtons per square centimeter (N), we used the following equations for resistance (R) and conductance (C). Note, our PDMS elastomer had 5 x 5 $mm^2$ dimensions, therefore to get the force in $N/cm^2$ we multiplied equation 3 by a factor of 4. The obtained calibration curve is shown below in SI Fig. 3.

$$R = \frac{(5000 - V_{out}) \cdot 10000}{V_{out}} \quad (1);$$

$$C = \frac{1000000}{R} \quad (2);$$

$$N = \frac{C}{80} \cdot 4 \quad (3)$$

**Functionality of traptavidin.** We made sure that our tAv was indeed functional and bound biotinylated 5 kb long DNA molecules. For that purpose we acquired TIRF images at 488 nm wavelength excitation and they showed no DNA binding on the surface modified by only m-PEG (SI Fig. 7A), while the m-PEG/bt-PEG mixture modified surface was densely covered with DNA molecules (SI Fig. 7B). Both surfaces were incubated with 0.02 mg/mL tAv, washed, incubated with 5 kb long DNA, washed and then stained with SG. These results verified functionality of the surface bound tAv molecules.

**Functionality of biotinylated antibodies directed against digoxigenin.** We prepared biotinylated anti-dig antibodies by conjugating them with Biotin-PEG4-NHS ester. Functionality of these bt-anti-dig conjugates was tested on a surface. The bt-anti-dig were immobilized on a PEGylated surface (10% bt-PEG) covered with sAv. Next, the singe-end dig-labeled λ DNA (dig-λ DNA) molecules were immobilized, stained using SG, and visualized using TIRF microscopy at 488 nm excitation without (SI Fig. 10A) or with buffer flow (SI Fig. 10B). These images showed mainly one-end tethered DNA fragments, suggesting successful anti-dig and dig-λDNA interaction. No binding of dig-λ DNA was observed on the PEGylated surface (10% bt-PEG) covered with only sAv.

**SI Table 1:** List of parameters of Si masters used in this work.

| Name of Si master | Line-width [nm] | Line-depth [nm] | Size of the patterned area [$cm^2$] | Comments |
|---|---|---|---|---|
| 1 | ~1000 | ~320 | 2.5 x 1.2 | Variable line-spacing ranging from 9.4 to 14.4 µm. |
| 2 | ~430 | ~200 | 0.7 x 0.8 | Variable line-spacing ranging from 12 to 14.5 µm. |
| 3 | ~180; ~900 | ~200 | 1.3 x 1.1 | Alternating line-width: first line thin and the following thick. Constant line-spacing of 12 µm. |
| 4 | ~500 | ~200 | 0.5 x 0.9 | Constant line-spacing of 12 µm. |
| 5 | ~1000 | ~200 | 0.5 x 0.9 | Constant line-spacing of 12 µm. |
| 6 | ~350 | ~200 | 0.5 x 0.9 | Constant line-spacing of 12 µm. |
| 7 | ~500 | ~200 | 0.5 x 0.9 | Constant line-spacing of 11 µm. |
| 8 | ~500 | ~200 | 0.5 x 0.9 | Constant line-spacing of 13 µm. |
| 9 | ~500 | ~200 | 0.5 x 0.9 | Constant line-spacing of 14 µm. |

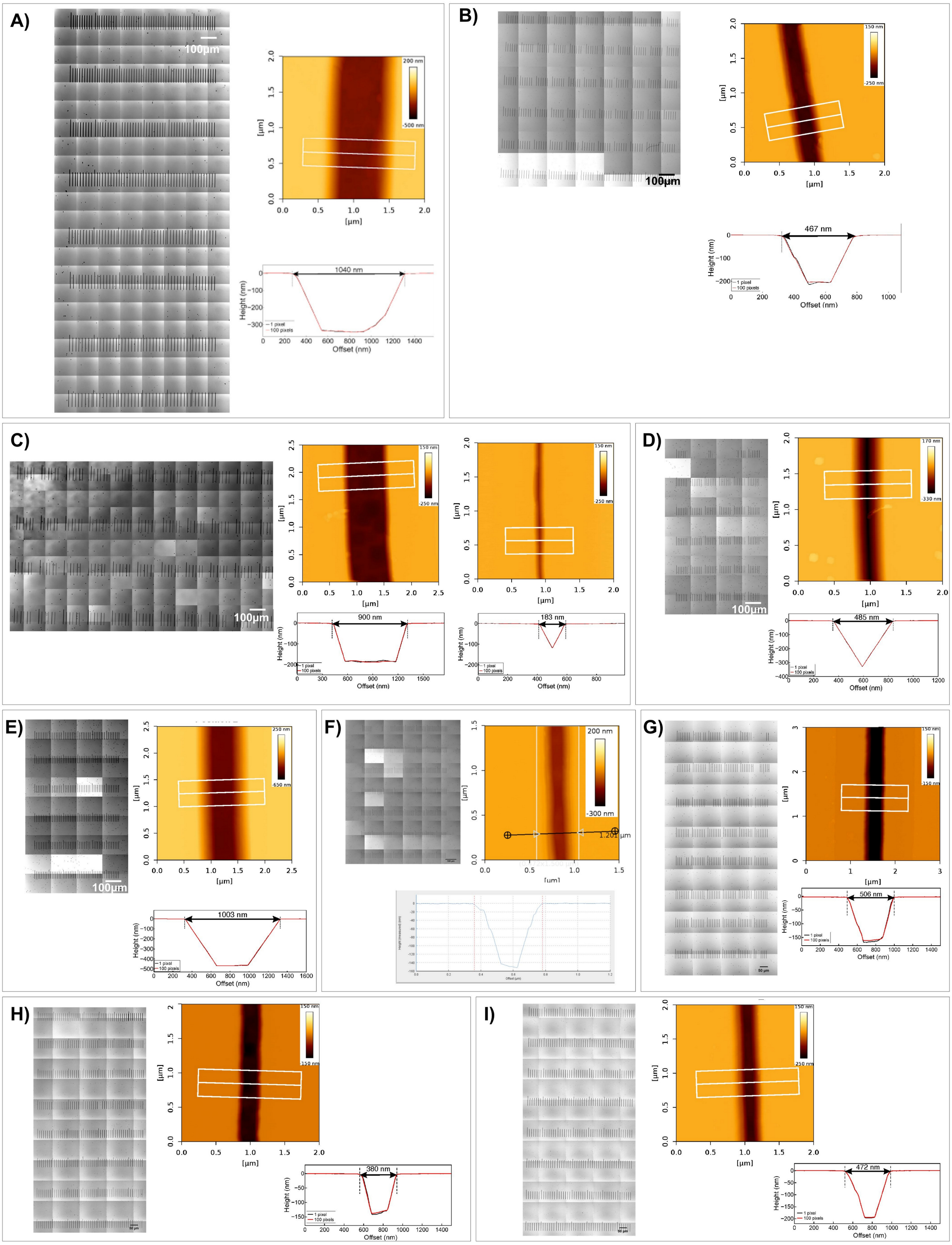


**SI Figure 1:** Overview of Si masters used in this work. **A – I )** On the left of each panel there is an optical image of the Si master #1 - #9 respectively after gold layer removal, on the middle there is an AFM image showing inscribed line profile and on the right bottom there are line-profiles (1 and 100 pix.) taken on the AFM image. For panel **C)** only difference is that this Si master contains alternating thick (~900 µm) and thin (~180 µm) lines.

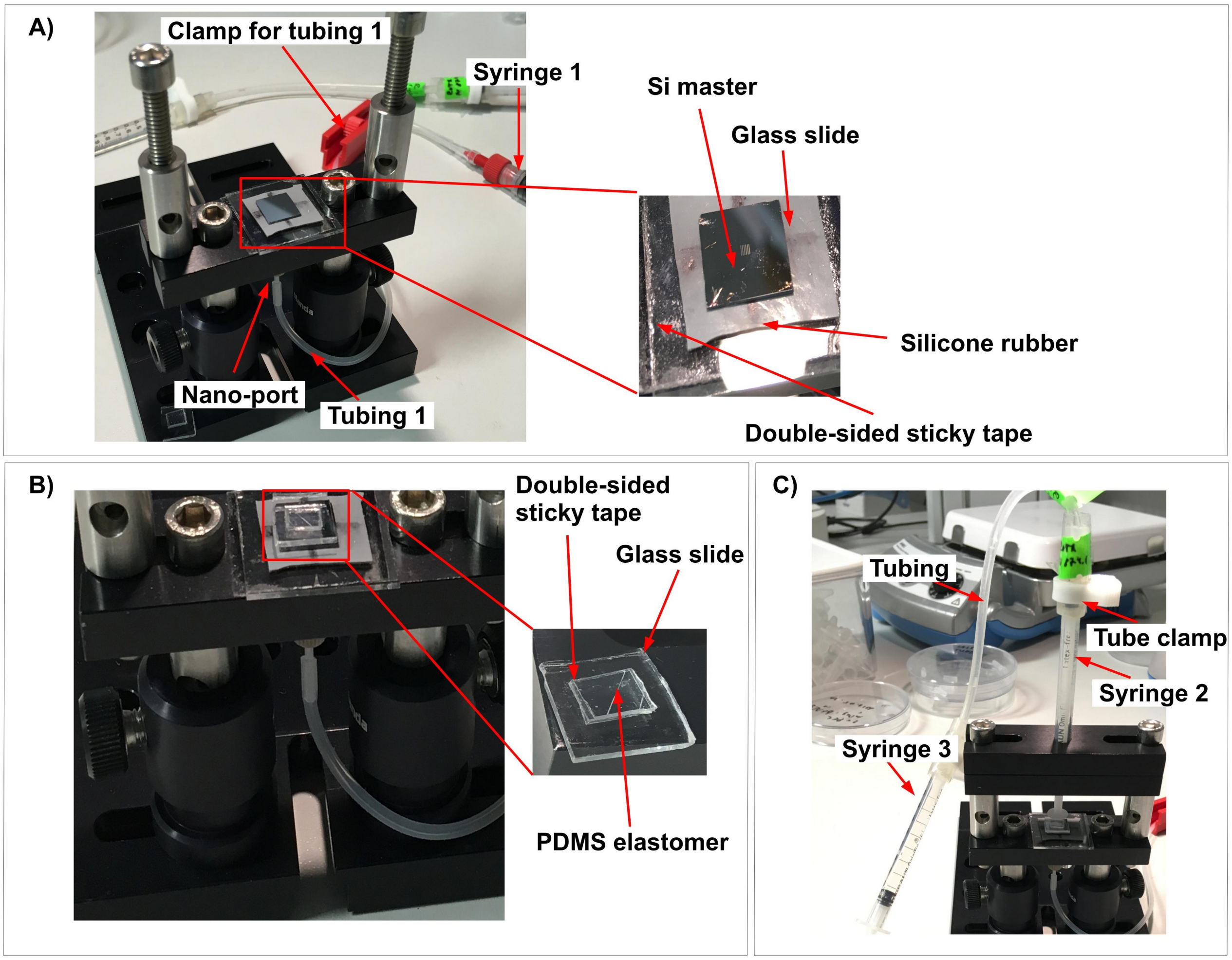


**SI Figure 2:** A portable printing device (PPD). **A)** Photograph of the PPD device with a Si master. **B)** The glass slide with attached PDMS elastomer was placed topside down onto the Si master manually using a tweezer. **C)** The system of two tubing-connected syringes was positioned above the glass slide with the PDMS elastomer, and printing pressure was applied by pressing the distant syringe (syringe 3). We controlled the printing pressure by monitoring the volume scale on the syringe 3.

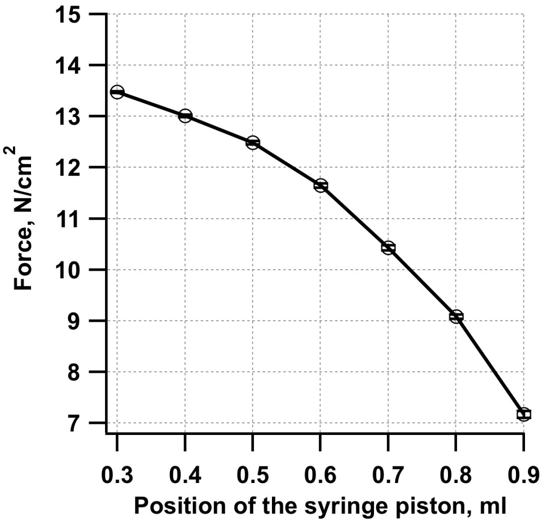


**SI Figure 3:** Syringe piston position and the printing force calibration curve of the portable printing device (PPD). Mean values (open circles) and SEM (error bars) represents six measurements.

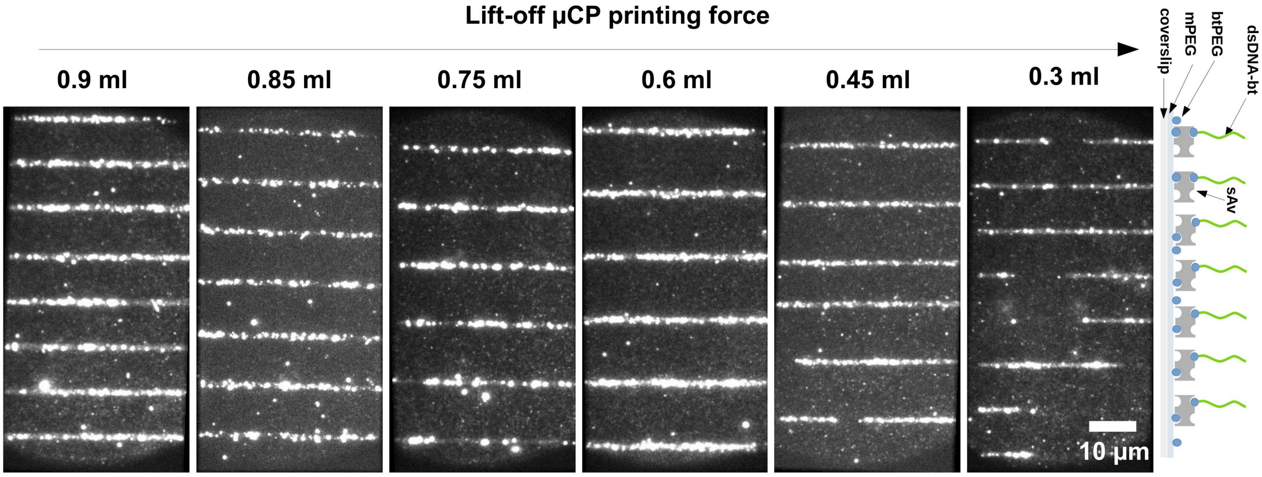


**SI Figure 4:** Characterization of short DNA molecule arrays and printing pressure effect on their quality. TIRF images of 5 kb long biotinylated DNA molecules stained with SYTOX green (SG), which were immobilized on the streptavidin (sAv) line-features fabricated on modified coverslip. Cartoon illustration on the right side depicts the immobilization scheme. Printing pressure, which was varied from the low (0.9 mL) to the relatively high (0.3 mL), is indicated above each image. [sAv] = 0.02 mg/mL, Si master #1, images presented here were acquired at 488 nm wavelength excitation and averaged over 10 consecutive frames.

**SI Table 2:** List of the quality factors (QF) obtained by changing the printing pressure (PP) of the lift-off µCP or streptavidin (sAv) and traptavidin (tAv) ink concentration during PDMS elastomer inking while keeping the PP constant. SEM – standard error of the mean.

| PP [mL] at 0.02 mg/mL of sAv | 〈QF〉 ± SEM | sAv concentration [mg/mL] at 0.6 mL PP | 〈QF〉 ± SEM | tAv concentration [mg/mL] at 0.6 mL PP | 〈QF〉 ± SEM |
|---|---|---|---|---|---|
| 0.9 | 12 ± 2.2 | 0.013 | 14.22 ± 2.1 | 0.06 | 6.5 ± 1.03 |
| 0.85 | 13 ± 1.7 | 0.017 | 21.1 ± 3 | 0.04 | 5.1 ± 0.3 |
| 0.75 | 16.8 ± 3.5 | 0.027 | 12.7 ± 1.1 | 0.03 | 10.4 ± 0.4 |
| 0.6 | 17.7 ± 2.7 | | | 0.02 | 10.3 ± 0.8 |
| 0.45 | 14.7 ± 3.1 | | | 0.015 | 5.7 ± 0.9 |
| 0.3 | 5.8 ± 0.7 | | | | |

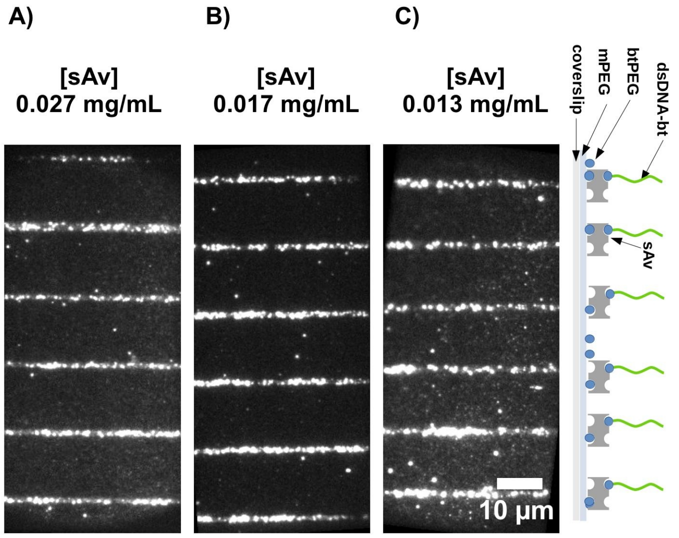


**SI Figure 5:** Characterization of short DNA molecule arrays and streptavidin (sAv) ink concentration effect on their quality at constant printing pressure (PP). **A-C)** TIRF images of 5 kb long biotinylated DNA molecules stained with SG, which were immobilized on the sAv line features fabricated on modified glass coverslip. Concentration of sAv ink: **A)** 0.027 mg/mL, **B)** 0.017 mg/mL, **C)** 0.013 mg/mL. Cartoon illustration on the right side depicts the immobilization scheme. PP = 0.6 mL, Si master #1, TIRF images were acquired at 488 nm wavelength excitation and averaged over 10 consecutive frames.

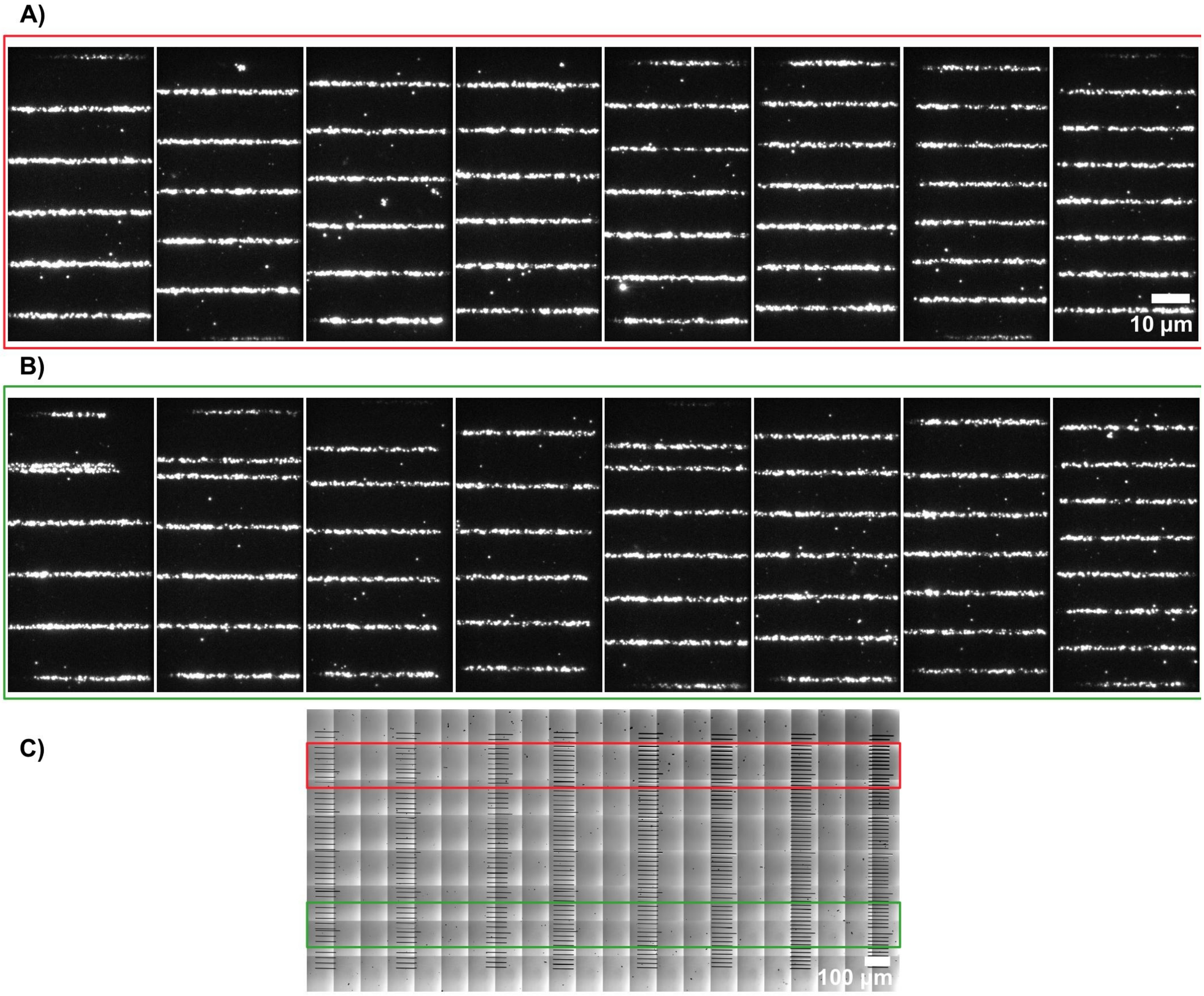


**SI Figure 6:** Printing quality across the full area of the template. **A-B)** TIRF micrographs of the single-tethered and SYTOX Green stained 5 kb DNA molecules immobilized on streptavidin template fabricated using Si master #1 and printing pressure (PP) equal to 0.6 mL. Excitation was at 488 nm wavelength and images are averages of 10 consecutive frames. **C)** Optical image of the Si master #1. Red and green squares represents region of interests for the **A** and **B** panel images.

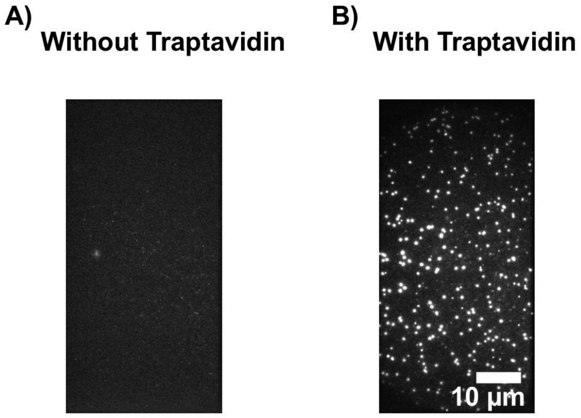


**SI Figure 7:** Functionality of surface immobilized traptavidin (tAv). The biotinylated 5 kb DNA molecules were immobilized on either **A)** bare or **B)** tAv covered (at 0.02 mg/mL) PEGylated (10% biotin-PEG) glass coverslip surface. DNA molecules were stained with the SYTOX Green, TIRF images represented here were acquired at 488 nm wavelength excitation and averaged over 10 consecutive frames.

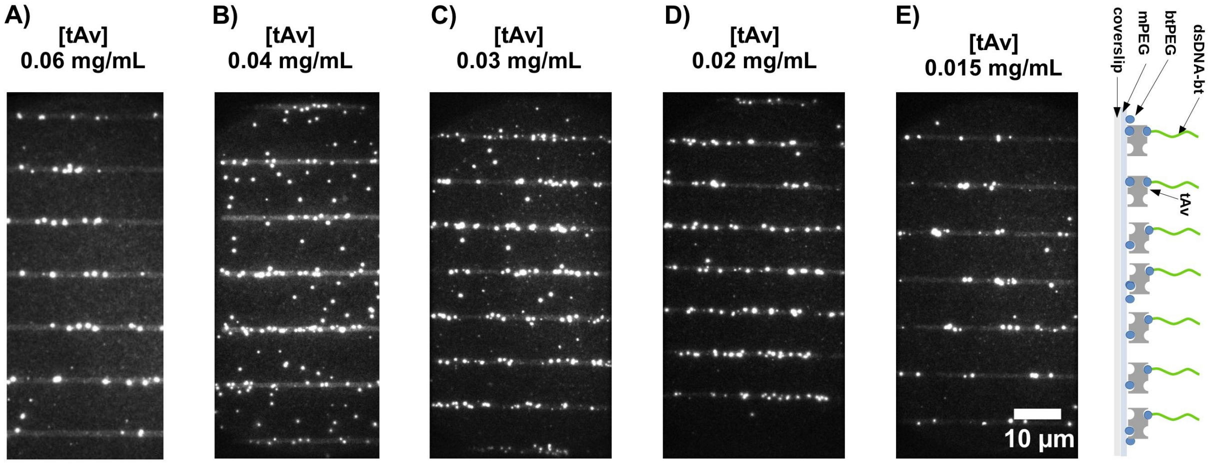


**SI Figure 8:** Characterization of short DNA molecule arrays and traptavidin (tAv) ink concentration effect on their quality at constant printing pressure (PP). **A-E)** TIRF images of 5 kb long biotinylated DNA molecules stained with SYTOX green (SG), which were immobilized on the tAv line features fabricated on modified glass coverslip. Concentration of tAv ink: **A)** 0.06 mg/mL, **B)** 0.04 mg/mL, **C)** 0.03 mg/mL, **D)** 0.02 mg/mL, **E)** 0.015 mg/mL. Cartoon illustration on the right side of the panel **E** depicts the immobilization scheme. PP = 0.6 mL, Si master #1, TIRF images were acquired at 488 nm wavelength excitation and averaged over 10 consecutive frames.

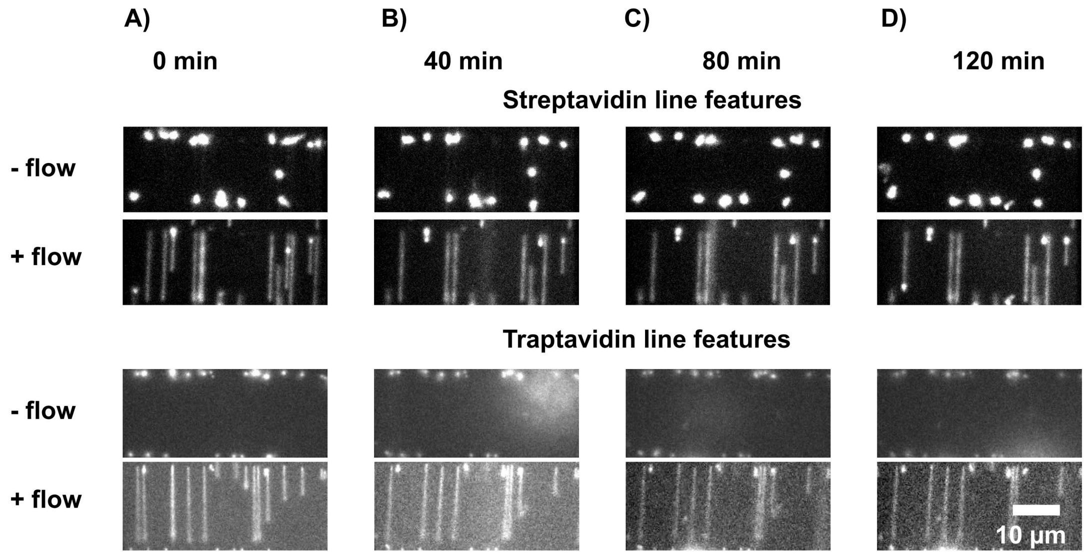


**SI Figure 9:** Stability test of single-tethered Soft DNA Curtains. **A-D)** An example of single-tethered λ DNA molecules immobilized on a streptavidin (top panels) or traptavidin (bottom panels) array template, stained with SYTOX green (SG). Images were acquired every 20 min for a period of 2 h. In between acquisitions, there was no buffer flow applied. During the acquisition, 20 frames were acquired at buffer flow of 1 mL/min and 20 frames without the flow. E) Printing pressure was 0.6 mL, Si master #3, TIRF images were acquired at 488 nm wavelength excitation and averaged over 10 consecutive frames.

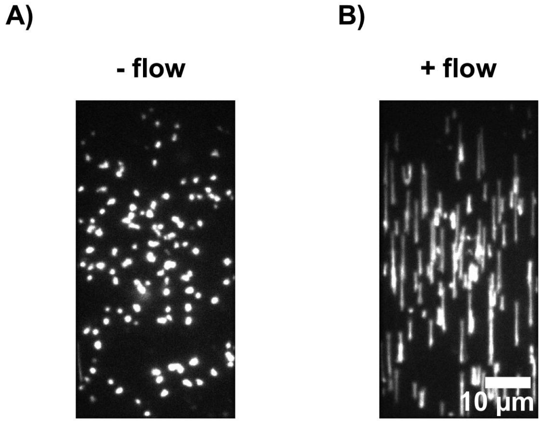


**SI Figure 10:** Functionality of surface immobilized biotinylated antibodies directed against dogoxigenin (bt-anti-dig). The digoxigenated λ DNA molecules were immobilized on PEGylated (10% biotin-PEG) glass coverslip surface via bt-anti-dig and streptavidin (sAv, at 0.02 mg/mL). TIRF Images of SYTOX Green stained DNA molecules were acquired **A)** without or **B)** with buffer flow. These images were acquired at 488 nm wavelength excitation and averaged over 10 consecutive frames.

**SI Table 3:** Dependency of anchoring efficiency of the second DNA end on line-width of line-features.

| Si master | Line-width of line-features (nm) | Separation distance between line-features (µm) | 〈% of both-end anchored DNA〉 ± SD | 〈# of both-end anchored DNA per line-feature〉 ± SD |
|---|---|---|---|---|
| #4 | 350 | 12 | 70.5 ± 34 (4.1 SEM, n=170) | 1.6 ± 1.2 (0.1 SEM) |
| #5 | 500 | 12 | 73.9 ± 23.1 (2.9 SEM, n=307) | 3.6 ± 1.8 (0.2 SEM) |
| #6 | 1000 | 12 | 78.6 ± 27.1 (3.5 SEM, n=380) | 4.9 ± 3.2 (0.4 SEM) |

**SI Table 4:** Dependency of anchoring efficiency of the second DNA end on line-separation distance of the tAv line-features.

| Si master | Line-width of line-features (nm) | Separation distance between line-features (µm) | 〈% of both-end anchored DNA〉 ± SD |
|---|---|---|---|
| #7 | 500 | 11 | 47.7 ± 28.8 (4.8 SEM, n=167) |
| #5 | 500 | 12 | 73.9 ± 23.1 (2.9 SEM, n=307) |
| #8 | 500 | 13 | 80 ± 16.7 (3 SEM, n=159) |
| #9 | 500 | 14 | 20.3 ± 22 (4 SEM, n=128) |

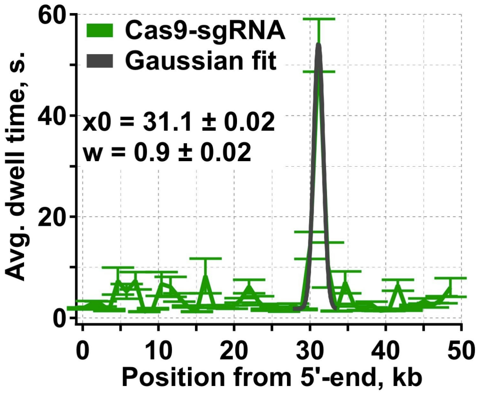


**SI Figure 11:** Histogram representing mean dwell time of Cas9-ATTO647N binding events. Data acquisition and the experimental scheme is explained in the Figure 3 of the main text.
